# Combination of the Systemin peptide with the beneficial fungus *Trichoderma afroharzianum* T22 improves plant defense responses against pests and diseases

**DOI:** 10.1101/2022.03.20.485010

**Authors:** Aprile Anna Maria, Coppola Mariangela, Turrà David, Vitale Stefania, Cascone Pasquale, Diretto Gianfranco, Fiore Alessia, Castaldi Valeria, Romanelli Alessadra, Avitabile Concetta, Guerrieri Emilio, Woo Sheridan Lois, Rao Rosa

## Abstract

*Trichoderma* spp. are among the most widely used plant beneficial fungi in agriculture. A novel approach to enhance their effectiveness in plant defense is to use the fungi in combination with bioactive molecules including plant-derived compounds. Here, we show that plant treatment with *Trichoderma afroharzianum* (strain T22) and Systemin (Sys), a tomato plant peptide active in triggering plant defense, confers protection against the fungal pathogens *Fusarium oxysporum, Botrytis cinerea* and the insect pest *Tuta absoluta.* The observed defensive response was associated with increased accumulation of metabolites and transcripts involved in the Jasmonic acid (JA) pathway. Our findings suggest that the innovative combination of *T. afroharzianum* T22 and Sys can result in a more effective and robust control of different biotic stress agents.

## 1. Introduction

Tomato (*Solanum lycopersicum* L.) is the second most important economic crop in the world after potato (*Solanum tuberosum* L.), with cultivation covering 4.85 million ha and fruit production totaling about 182.3 million tons every year (FAOSTAT, 2019). Regardless of the type of cultivation or the plant life-cycle, increasing amounts of pesticides are required to protect tomato plants from pathogens and pests worldwide (Almeida et al., 2019; Savary et al., 2019; S Abbas et al., 2020) with well-known undesirable effects on the environment (pollution, soil depletion, biodiversity loss) and human health (Rani et al., 2021). Public concerns prompted the research for alternative strategies of plant protection that are effective, reliable and sustainable (Iriti and Vitalini, 2020). In this context, biological control of plant diseases through the application of bioagents is a highly promising approach (Vinale et al., 2008). Among biological control agents (BCA), beneficial fungi belonging to *Trichoderma* spp. represent the most widespread soil microbes used in agriculture and a major constituent of commercially available biological products (i.e., biopesticides, biostimulants, biofertilizers) (Woo et al., 2014). Different *Trichoderma* species and their metabolites act against plant pathogens through several mechanisms, including direct antagonism of disease agents (mycoparasitism) (Poveda, 2021), antibiotic-mediated suppression, secretion of lytic enzymes and other byproducts, competition for nutrients (Al-Ani, 2018). Additionally, *Trichoderma* root colonization induces a state of alert (defense priming) in plants leading to a faster and stronger defense response against future pathogen attacks, this is achieved reducing the effector-triggered susceptibility (ETS) and concurrently enhancing the effector-triggered immunity (ETI) (Hermosa et al., 2012; Martínez‐Medina et al., 2017). Within this scenario, the use of inducers of plant endogenous defenses is considered as one of the most promising approaches for the promotion of sustainable agricultural practices that may reduce synthetic chemical inputs (Burketova et al., 2015).

A pioneering study showed that plant root colonization by *T. afroharzianum* is associated with a reduction of the symptoms produced by the necrotrophic fungus *B. cinerea* as a consequence of systemic defense responses elicitation (De Meyer et al., 1998). Corroborating these findings, other studies showed the ability of *Trichoderma* to induce a set of defense-related genes associated with the JA and ethylene (ET) pathways (Shoresh et al., 2005;Korolev et al., 2008). Notably, *Trichoderma* spp. do not only protect against microorganisms but also against different insect pests (Muvea et al., 2014; Coppola et al., 2017; Contreras-Cornejo et al., 2018). Coppola et al (Coppola et al., 2019a) demonstrated that *Trichoderma atroviride* induced defenses are effective against two tomato pests with distinct eating habits: the piercing–sucking aphid *Macrosiphum euphorbiae* and the chewing noctuid moth *Spodoptera littoralis*. Moreover, plant colonization by the fungus activated indirect defenses via the production of volatile organic compounds (VOCs) that attracted the pest parasitoid *Aphidius ervi.*

It is well known that plant protection mediated by *Trichoderma* spp. can be deeply influenced by environmental conditions and ecological fitness of biocontrol agents hence being poorly predictable *a priori* (Howell, 2003; Alabouvette et al., 2009; Lecomte et al., 2016; Malik et al., 2018). Consequently, the sole use of plant beneficial microorganism may not be always sufficient to fully protect plants against biotic stresses. To overcome such limitations, new approaches to increase plant resistance may focus on the combined application of different resistance inducers (*i.e.* chemical and/or biological) (Zehra et al., 2017; Siah et al., 2018).

Among plants-derived molecules, the tomato peptide Sys has proved to be a very efficient elicitor of plant defense in tomato, grapevine, and eggplant (Coppola et al., 2019c; Molisso et al., 2021). Sys was the first peptide signal to be discovered in plants (Pearce et al., 1991). It is an octadecapeptide proteolytically released, upon wounding and herbivore attack, from the carboxy terminus of its 200 amino-acid precursor ProSystemin (ProSys) (Ryan and Pearce, 2003). In plants, the autocrine interaction of Sys with its receptor is followed by the activation of the octadecanoid pathway and the production of JA and its derivatives (JAs) (Pearce et al., 1991). The role of Sys in the defense response against wounding or herbivory was evaluated in several studies exploiting transgenic plants overexpressing or downregulating ProSys. Its overexpression increased not only the resistance of plants against lepidopteran larvae, fungi and aphids, but also the tolerance to salt stress (McGuRL et al., 1994; Orsini et al., 2010; Coppola et al., 2015), while its downregulation strongly increased plant susceptibility to herbivores (Orozco-Cardenas et al., 1993).

Recently, it was demonstrated that exogenous Sys supply induces an array of defense-related responses leading to the final accumulation of proteinase inhibitors (PINs) and other molecules able to interfere with the colonization of fungal pathogens or the growth and survival of herbivores (Coppola et al., 2019c; Molisso et al., 2021). Also, Sys-treated plants proved to be more attractive for pest natural enemies as a consequence of the modified blend of volatile compounds emitted, which is known to be used by predators and parasitoids of phytophagous insects to locate their preys and hosts (Ninkovic et al., 2021).

Given that both *Trichoderma* and Sys activate the JA pathway, a key hub for plant defense and resistance towards insects and fungi, we hypothesized that their combined treatment on plants could result in an amplified disease and pest control outcome. Accordingly, here we show that their combination confers tomato resistance towards both the necrotrophic and hemibiotrophic fungal pathogens *B. cinerea* and *F. oxysporum,* respectively, and the insect pest *T. absoluta*. Our data provides evidence that plant treatments in a combination of beneficial fungi and plant derived resistance inducers might represent an effective tool for sustainable pest management.

## 2. Materials and methods

### 2.1 Preparation of fungal cultures and inoculum and insects rearing

Fungal cultures were obtained from the collection available at the Department of Agricultural Sciences of the University of Naples Federico II. *T. afroharzianum (*formerly*-T. harzianum)* strain T22 (T22), was isolated from the commercial product Trianum-P (Koppert Biological Systems, Rotterdam, the Netherlands), cultured on potato dextrose agar (PDA; HiMedia) and grown at 25°C until complete sporulation. Conidia were collected in sterile distilled water by scraping the surface of sporulating fungal cultures and recovering the spores by centrifugation at 2,700 g for 10 min. Conidia concentration was adjusted to 10^7^ spores/mL for subsequent bioassays.

*F. oxysporum* f.sp. *lycopersici* (*Fol*) race 2 isolate 4287, and the *Fol* knockout mutant *fmk1*Δ were grown in potato dextrose broth (PDB) at 28°C with orbital shaking at 170 rpm. Phleomycin (5.5μg/ml) was added to the culture medium when required. Fresh conidia were separated from the mycelium by filtering 5 day-old cultures through a nylon filter membrane (mesh size 10 μm) and collected after centrifugation at 2,700 g for 10 min. Storage of the obtained conidia was performed as previously described (Di Pietro et al., 2001).

*B. cinerea* was grown on MEP (Malt extract with peptone) solid medium at 22°C. Conidia were collected by washing the agar surface with sterile distilled water containing 0.1% Tween 20, filtered, and adjusted to a concentration of 1× 10^6^ conidia /ml.

The original strain of *T. absoluta* was collected in 2019 in tomato greenhouses located in Battipaglia (Salerno, Italy)(Gontijo et al., 2019). The insect was continuously reared at the National Research Council, Institute for Sustainable Plant Protection (CNR-IPSP, Portici), inside aerated cages (Vermandel ®, The Netherlands) at 22± 5%°C, 65± 5% relative humidity (RH) and 16L:8D photoperiod.

### 2.2 Peptide synthesis

The Sys peptide synthesis, and purification was obtained as previously described (Coppola et al., 2019c).

### 2.3 In Vitro Trichoderma-Sys compatibility assay

To assess if Sys had a direct antimicrobial effect on *T. afroharzianum T22*, an *in vitro* assay was carried out to measure fungal growth in presence or absence of different concentrations of the peptide. Antifungal assays were conducted as previously described (Pastor-Fernández et al., 2020). Wells of a sterile 12-well plate were filled with 1 ml of half-strength potato dextrose broth (PDB 1/2) medium containing Sys, at a final concentration of 100 pM or 100 fM except for the negative control wells. The positive control was filled with 200 μg/ml Switch^®^ fungicide (Syngenta, 37.5% w/w cyprodinil and 25% w/w fludioxonil). Conidia of *T. afroharzianum* T22 were then added to each well to reach a final concentration of 10^4^ conidia/ml. Each plate was then placed in a shaker and incubated for 24 hours at 25 ± 1°C. End-point fungal growth was assessed by measuring the medium turbidity at a wavelength of 600 nm (OD_600_) on BioPhotometer Spectrophotometer UV/VIS (Eppendorf, Hamburg, Germany).

### 2.4 Seed Treatment and Plant Growth

Seeds of *Solanum lycopersicum* cultivar “San Marzano Nano” were surface sterilized with 2% sodium hypochlorite for 10 min, then fully rinsed in sterile distilled water. For the seed treatments, washed tomato seeds were either coated by immersion in a fresh spore suspension (final concentration of 10^7^ conidia/mL) of *T. afroharzianum* T22, or treated with water (Ctrl), stirred frequently to uniformly cover the seed surface, left to air dry for 24 h and stored at 4^◦^C until use. Seeds were germinated on Whatman^®^ sterile filter paper (Sigma-Aldrich, Darmstadt, Germany), moistened with sterile distilled water and placed in a growth chamber at 24 ± 1°C and 60 ± 5% RH in the dark. At the emergence of the cotyledons, germinated seedlings were individually transplanted into pots containing sterile commercial soil (Universal Potting Soil, Floragard, Oldenburg, Germany) and kept in a growth chamber at 26 ± 1°C and 60 ± 5% RH., 18L:6D-photoperiod by lamps of 5,000 lux. After 2 weeks, tomato seedlings were transplanted into 10-cm diameter plastic pots containing sterilized soil and grown for 2 weeks under the same environmental conditions. For *F. oxysporum* assay, seeds were planted on wet vermiculite in plastic trays and incubated in plant growth chambers for two weeks at 26 ± 1°C and 60 ± 5% RH.

### 2.5 In vivo F. oxysporum infection assays

Two weeks-old plants (Ctrl and T22 seed-treated), treated with Sys or PBS buffer 1X dissolved in 5 ml of distilled water (final Sys concentration 100 fM), were tested for resistance to *F. oxysporum.* Twenty four hours after Sys application *Fol* infection was performed by dipping tomato roots in a microconidia suspension (5×10^6^ conidia/ml) from *Fol* wt or *fmk1Δ* strains (Di Pietro and Roncero, 1998). Tomato seedlings were planted into minipots containing vermiculite and maintained in a growth chamber (28 °C; 14L:10D photoperiod). Plant infection was recorded daily up to 20 days and percentage of plant survival calculated by the Kaplan–Meier method (López-Berges, 2012). Fifteen plants per treatment were used and each experiment was repeated in three independent occasions.

### 2.6 In vivo B. cinerea infection assays

The application of Sys and *T. afroharzianum* was tested either alone or in combination to evaluate the effects of each treatment on *B. cinerea* leaf colonization. Fully expanded leaves from four week-old plants (Ctrl and T22) were treated with 15 spots of 2 μL of 100 fM Sys or PBS; spots were produced on the abaxial surface using a pipette. Leaves were tested for resistance to *B. cinerea* after six hour*s.* by applying 10 μl drops of *B. cinerea* spores (1×10^6^ conidia/ml) between tomato leaf veins, using 8 different inoculation points per leaf. from 5 different plants for each individual treatment. Detached leaves were placed on sponges soaked in sterile water and incubated in a growth chamber at 23°C, under 16L:8D photoperiod and 90% ± 5% RH. Disease severity was quantified by measuring the development of necrotic leaf areas caused by the fungal pathogen over time at different days post inoculums (DPI) using a digital caliper (Neiko 01407A; Neiko Tools, Taiwan, China).

### 2.7 In vivo T. absoluta assays

To evaluate the possible induction of repellency linked to the alteration of the profile of volatile organic compounds released, a choice test was performed using a random block experimental design. Three 4-weeks old plants (Ctrl and T22) were treated with 100 fM Sys or PBS by foliar spotting. After 6 hours, plants were placed into a single mesh cage (60×60×180 cm; Vermandel, The Netherlands) at a distance of 20 cm from each other. Three to 5 days old mated females of *T. absoluta* were released into the cage keeping a 1:1 ratio between them and the plants. After 2 days (oviposition period), females were removed from the cage by an insect aspirator and the eggs laid were counted. The test was replicated four times totalling twelve females/plant tested. To evaluate antibiosis/antixenosis, a no-choice test was performed by feeding *T. absoluta* larvae with detached leaves from each treatment and relative control. Fully expanded leaves to be used in the test were cut from plants treated as described before and immediately placed through the petiole in a 5-mL tube capped by cotton wick, sealed by parafilm tape and filled with tap water. Larvae to be used in the test came from tomato plants exposed for 24 h to hundreds of *T. absoluta* adults in a mesh cage. Tomato leaves with eggs were transferred into aerated cages and a single hatched larva was gently transferred onto a detached leaf by a soft brush and kept into a plastic box (Duchefa, The Netherlands) constituting the experimental unit. Thirty larvae were tested for each treatment. Leaves were examined daily to check the larval longevity and survival and to record the time to pupation.

### 2.8 Gene expression analysis

Expression levels of defense-related genes in control and *T. afroharzianum* - and/or Sys-treated tomato plants were quantified by real-time PCR (RT-PCR). The selected genes were: *Allene Oxide Synthase* (*AOS;* Solyc11g069800), *Lipoxygenase D* (*LoxD;* Solyc03g122340), *Wound-induced Proteinase Inhinbitor I* (*Pin I;* Solyc09g084470), *Threonine deaminase* (*TD;* Solyc09g008670), and *Leucine aminopeptidase* (*LapA;* Solyc12g010030). Fully expanded leaves from 4 weeks old plants (Ctrl or T22) were collected 6 hours after Sys treatment and immediately frozen in liquid nitrogen.

Total RNA extraction, first strand cDNA synthesis and RT-PCR were performed according to standard procedures, as reported elsewhere (Corrado et al., 2012). RT-PCR was performed using Rotor Gene 6000 (Corbett Research; Sydney, Australia). For each sample, two technical replicates from each biological replicate were used for gene expression analysis. The housekeeping gene EF-1α was used as endogenous reference gene for the normalization of the expression level of target genes (Müller et al., 2015). Relative quantification of gene expression was carried out using the 2^−ΔΔCt^ method and referred to the mock-treated control (seeds treated with PBS 1X) (relative quantification, RQ = 1). Primers and their main features are reported in Additional file 2: Table S1.

### 2.9 LC-HESI(−)-HRMS of JA and JA-related metabolites

By using LC-HESI(−)-HRMS analysis, the levels of JA, its glycosylated (tuberonic acid-glucoside) and methylated form (MeJA) as well as three of its intermediates (13(S)-HPOT, 12,13(S)-EOT and 12-OPDA) were measured in control plants or in plants treated with Sys, T22 or both. Detection and quantification of JA and JA-related metabolites was performed as previously reported with slight modifications (Liu et al., 2010;Van Meulebroek et al., 2012;Grosso et al., 2018). Briefly, 100 mg frozen leaf samples of 4 weeks old plants (Ctrl or T22) were collected 6 hours after Sys treatment and homogenized with 1ml cold extraction buffer (methanol, ultrapure water and formic acid (75:20:5, v/v/v) spiked with 10 μg ml-1 formononetin as internal standard), vortexed continuously at −20°C o.n. Following centrifugation at 15,000 g for 20’ at 4°C, 500 ml were transferred to 30 kDa Amicon^®^ Ultra centrifugal filter unit (Merck Millipore Corporation, Massachusetts, USA), and centrifuged for 15’ at 15,000 g and 4 ◦C. Finally, extracts were evaporated by speedvac until a final volume of 50 μl was reached, and then transferred to the LC vials. LC-HESI(−)-HRMS analyses were carried out using a Dionex high performance liquid chromatography-diode-array detector (LC-DAD) coupled to high resolution mass spectrometry equipped with a heated electrospray source operating in negative ione mode (ESI(−)HRSMS) (Thermo Fisher Scientific, Waltham, MA, USA). LC analysis was performed as reported before (Liu et al., 2010; Van Meulebroek et al., 2012; Grosso et al., 2018), whereas mass spectrometry analysis was performed using a quadrupole-Orbitrap Q-exactive system (Thermo Fisher scientific, USA), using a single ion monitoring (SIM) method. More in detail, metabolite ionization was carried out using the following parameters: nitrogen was used as sheath and auxiliary gas (40 and 15 units, respectively); capillary and vaporizer temperatures set at 300°C and 280°C, respectively, discharge current at 3.5 KV, probe heater temperature at 360 °C, S-lens RF level at 50 V. The acquisition was carried out in the 110/1600 m/z scan range, according to the following parameters: resolution 70,000, microscan 1, AGC target 1e6, and maximum injection time 50. Data were analyzed using the Xcalibur 3.1 software (ThermoFisher scientific, USA). Metabolites were identified based on their accurate monoisotopic and adduct masses (m/z) and MS fragmentation, using both in house database and public sources (i.e. KEGG, MetaCyc, ChemSpider, PubChem, Metlin, Phenol-Explorer), as well by comparing chromatographic and spectral properties with authentic standards, when available. Relative abundances of the metabolites studied were calculated using the Xcalibur 3.1 software (Thermo Fisher scientific, USA). JA, and JA-related metabolites were quantified in a relative way by normalization on the internal standard amounts. Data are presented as means and standard deviations of three independent biological replicates.

### 2.10 Statistical analysis

Differences in the survival rate of tomato plants infected by *F. oxysporum* and tomato plants infested by *T. absoluta* larvae were analyzed using the Kaplan-Meier method and compared among groups using the log-rank test. One-Way ANOVA test, followed by Tukey’s post-hoc multiple comparison test (P< 0.05), were used to evaluate: differences in relative transcripts abundance, in necrosis diameter development, the effect of Sys on T22 growth, and in *T. absoluta* oviposition rate. Welch’s ANOVA test followed by Dunnett’s T3 test was used to evaluate differences *T. absoluta* larval longevity. Student’s t-test was used to identify metabolites whose abundance significantly changed with respect to the control condition

## 3. Results

### 3.1 Sys peptide did not directly impact T. afroharzianum T22 growth

Sys treatment did not impact on T22 proving that Sys does not have inhibitory effect on fungal growth (Additional file 1: Figure S1).

### 3.2 Sys and T. afroharzianum T22 reduce F. oxysporum infection

To evaluate if treatments with Sys and/or *T. afroharzianum* T22 were able to control *F. oxysporum* wilt disease on tomato plants, roots of two-week-old seedlings were left uninoculated, inoculated with the *Fol* wt strain or with the *Fol fmk1*Δ avirulent mutant strain and survival of treated versus untreated plants was recorded over time. As expected, in all experiments and regardless of the applied treatment (PBS control, Sys *or* T22), uninoculated or *fmk1*Δ inoculated plants displayed neither wilting symptoms nor mortality throughout the whole evaluation process (Additional file 1: Figure S2).

On the contrary, most of tomato plants germinated from PBS soaked seeds (Ctrl) and inoculated with the *Fol* wt strain showed progressive wilting symptoms and died after 19 days post inoculation (Additional file 1: Figure S2). Seed treatment with Sys or coated with *T. afroharzianum* T22 only partially, though not significantly, decreased *Fol*-mediated wilting and death of plants (Figure 1A, B) (T22+Fol, Log-Rank test, χ^2^ = 1.588, df = 1; P = 0,2077; Sys+ Fol, Log-Rank test, χ2 = 0.771, df = 1, P = 0,3797).

**Figure 1.**
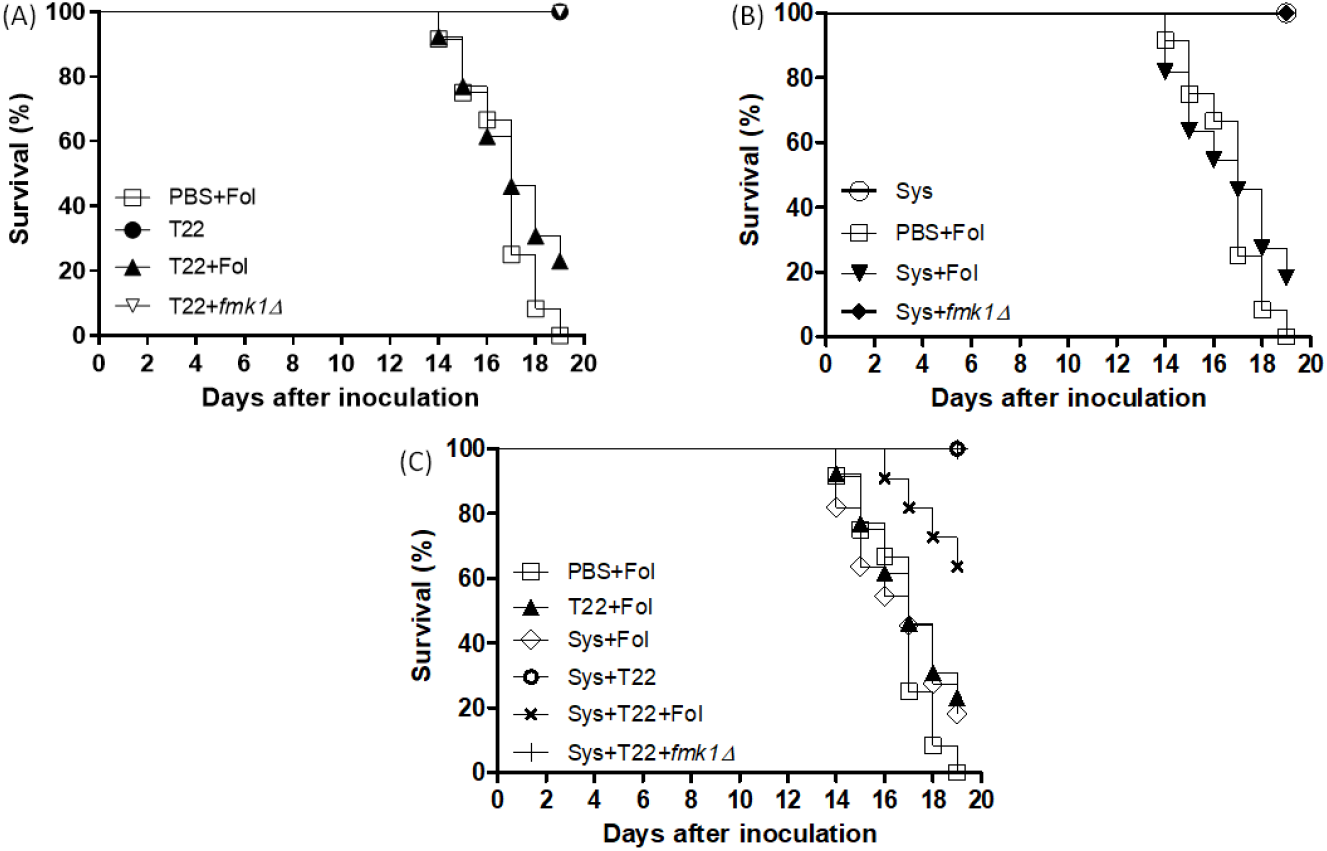
*T. afroharzianum* and Sys additively reduce *F. oxyporum* virulence on tomato plants. Kaplan– Meier plots showing survival of tomato plants treated with PBS (**A**, **B**, **C**), *T. afroharzianum* (**A**), Sys (**B**), or both (**C**), and not inoculated or inoculated by dipping roots into a suspension of 5×10^6^ conidia/ml of the indicated *Fol* strains. Experiments were performed at least three times with similar results. Percentage survival of tomato plants was plotted for 20 days. Data shown are from one representative experiment.

Noteworthy, combined treatment of tomato seeds with both Sys and *T. afroharzianum* T22 showed a stronger and significant reduction of plant mortality (Sys+T22+Fol, Log-Rank test, χ^2^ = 12.28, df = 1, P= 0,0005) as only 40% of these plants had died after 19 DPI, contrarily to those derived from PBS soaked seeds (Ctrl) where 100% mortality was observed (Figure 1C).

### 3.3 Sys and T. afroharzianum reduce B. cinearea infection

Plants treated with Sys alone or in combination with T22 showed a significant reduction of foliar damage already one day after *B. cinerea* inoculation. Conversely, T22 alone treated plants exhibited significant differences in disease severity only after 5 days post inoculation when compared to control plants. A further decrease in necrotic leaf area development could be observed 7 DPI in plants concomitantly treated with both Sys and T22, with about 63% reduction in disease severity compared to the control. Disease severity in these plants was significantly lower than the one observed in plants where each treatment was applied separately (− 52 % for Sys treatment and − 40% for T22) (Figure 2).

**Figure 2.**
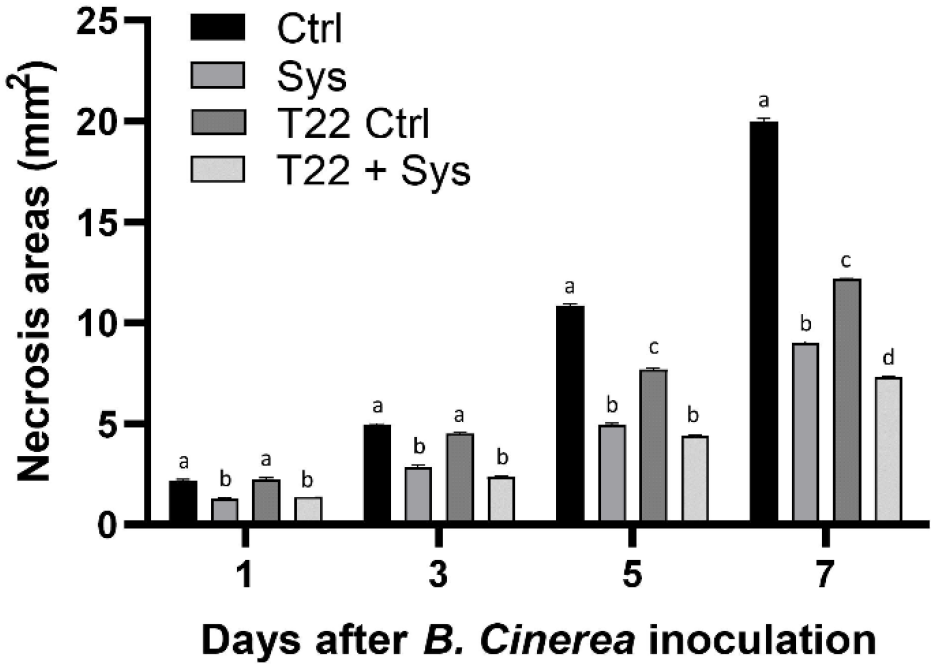
*T. afroharzianum* and Sys reduce *B. cinerea* virulence on tomato plants. Necrotic areas observed on *B. cinerea* inoculated leaves from tomato plants germinated from uncoated (Ctrl) or *T. afroharzianum* - coated seeds and treated or not with 100 fM Sys. The graph displays the average area of necrotic lesions formed 1, 3, 5, and 7 DPI. Letters indicate different statistical groups (One-Way ANOVA, P<0.05). Error bars indicate standard error (n = 90).

### 3.4 Sys exogenous supply and *T. afroharzianum* colonization alters *T. absoluta* infestation

The number of eggs laid per plant was influenced by treatments (Figure 3A, ANOVA, df=3, F=10.03, P<0.001) resulting lowered by Sys and raised by T22 colonisation. The combined application of T22 and Sys didn’t affect this parameter. Larvae survival curves revealed a significant antibiotic effects of *T. afroharzianum* inoculation and Sys treatment to *T. absoluta* development when used alone or in combination (Figure 3B, Log-Rank test, χ^2^ = 21.6, df = 3, P < 0.001). Survival rates, from eggs to pupation, were significantly lower for larvae fed on T22-treated leaves (Log-Rank test, χ^2^ = 7, df = 1, P = 0.008) and for larvae fed on Sys-treated leaves (Log-Rank test, χ^2^ = 18, df = 1, P < 0.001), in respect to control plants (Figure 3C).An additive effect was recoded for combined T22-Sys treatment (Pairwise log-Rank test T22 vs T22+Sys χ^2^ =10.64, df = 3, P = 0.014). Similarly, larval longevity was lowered by the combined application of T22 and Sys while no significant effect was recorded when the treatments were tested alone (Figure 3B, Welch ANOVA, F_3,146_=2.708, P =0.047).

**Figure 3.**
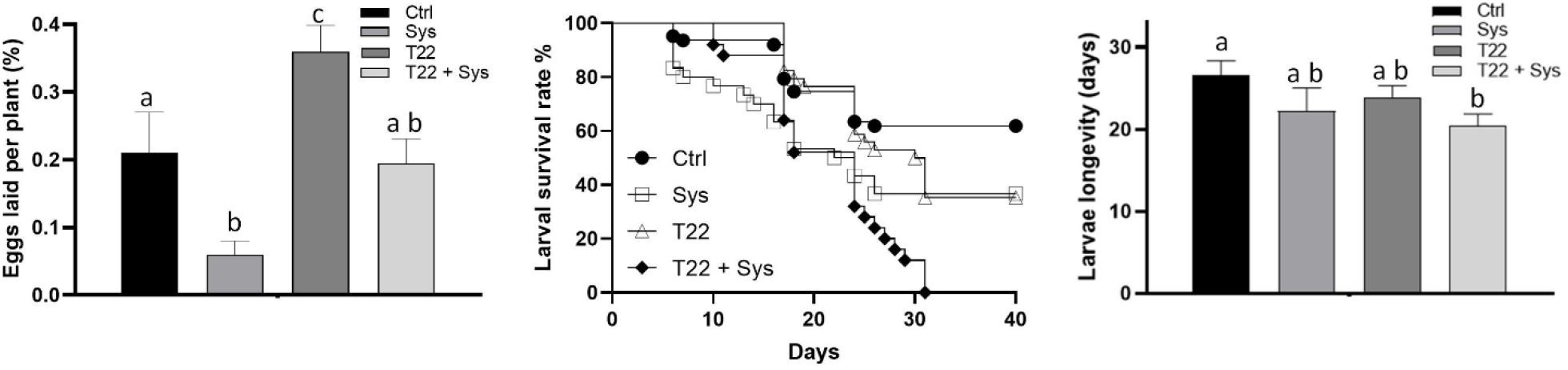
T. absoluta assays. (A) Larvae survival curves and (B) larvae longevity reared on tomato plants treated with Sys and *T. afroharzianum* T22, their combinations and relative control. (C) Eggs laid per plant by *T. absoluta* adults on the tomato plants treated as above described.

### 3.5 Sys and T. afroharzianum enhance the expression of plant defense-related genes

A significant increase of *AOS, Pin I, LapA* and *TD*, but not of *LoxD*, transcripts was recorded after Sys or T22 application (Figure 4). All gene transcripts, including those of the *LoxD* gene, were significantly more expressed in plants simultaneously treated with Sys and T22 compared to those treated with Sys or T22 alone.

**Figure 4.**
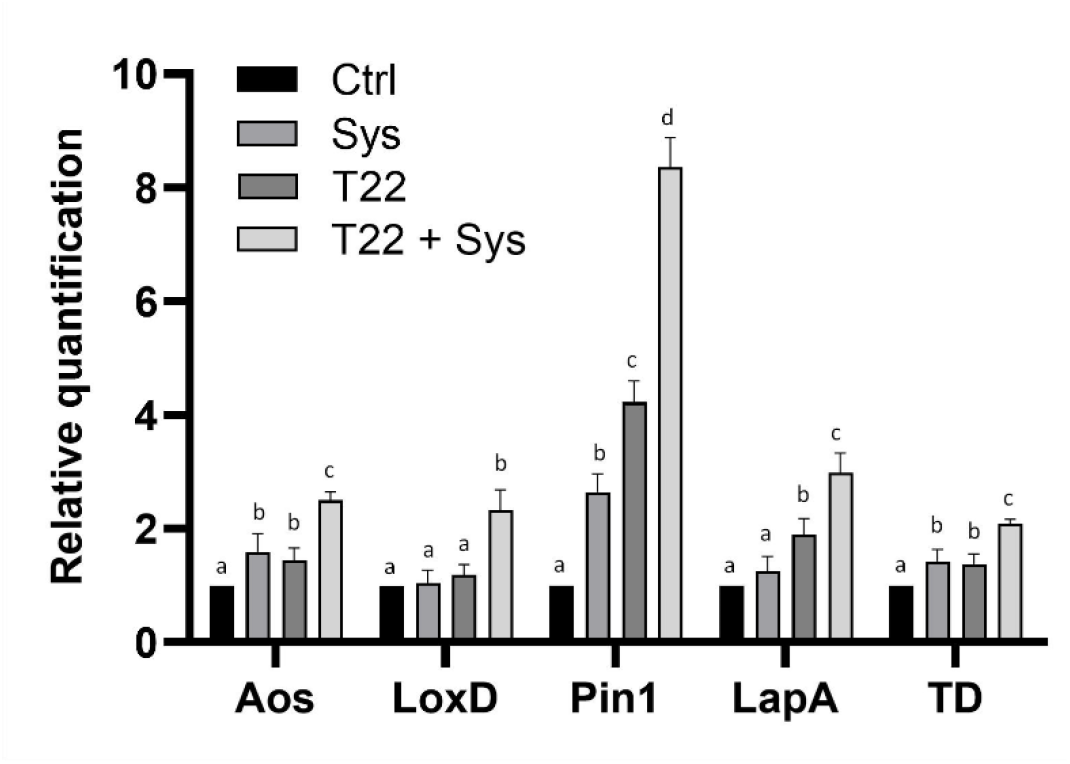
Level of defense-related genes expression in tomato plants grown from uncoated (Ctrl) or *T. afroharzianum* −coated seeds (T22) and treated or not with 100 fM Sys. Transcript levels of the following genes were analyzed: *Allene Oxide Synthase* (*AOS*), *LipoxygenaseD* (*LoxD*), *Wound-induced Proteinase Inhibitor I* (*Pin I*), **Threonine deaminase (TD), and** *Leucine aminopeptidase A* (*LapA*). Quantities of transcripts are relative to the calibrator control condition, plants grown from seeds that had been treated with PBS 1X (Ctrl). Different letters indicate statistically significant differences (One-Way ANOVA, P<0.05). Error bars indicate standard error (n=3).

### 3.6 Sys exogenous supply and T. afroharzianum colonization increase the accumulation of metabolites involved in the JA pathway

JA over-accumulated in plants treated with Sys and T22, alone or in combination, with a maximum fold of 1.74 in T22+Sys samples (Figure 5); in addition, Sys treated plants also displayed higher levels in 12-OPDA in respect to controls, while another intermediate, 12,13(S)-EOT, resulted more highly accumulated in T22+Sys plants.

**Figure 5.**
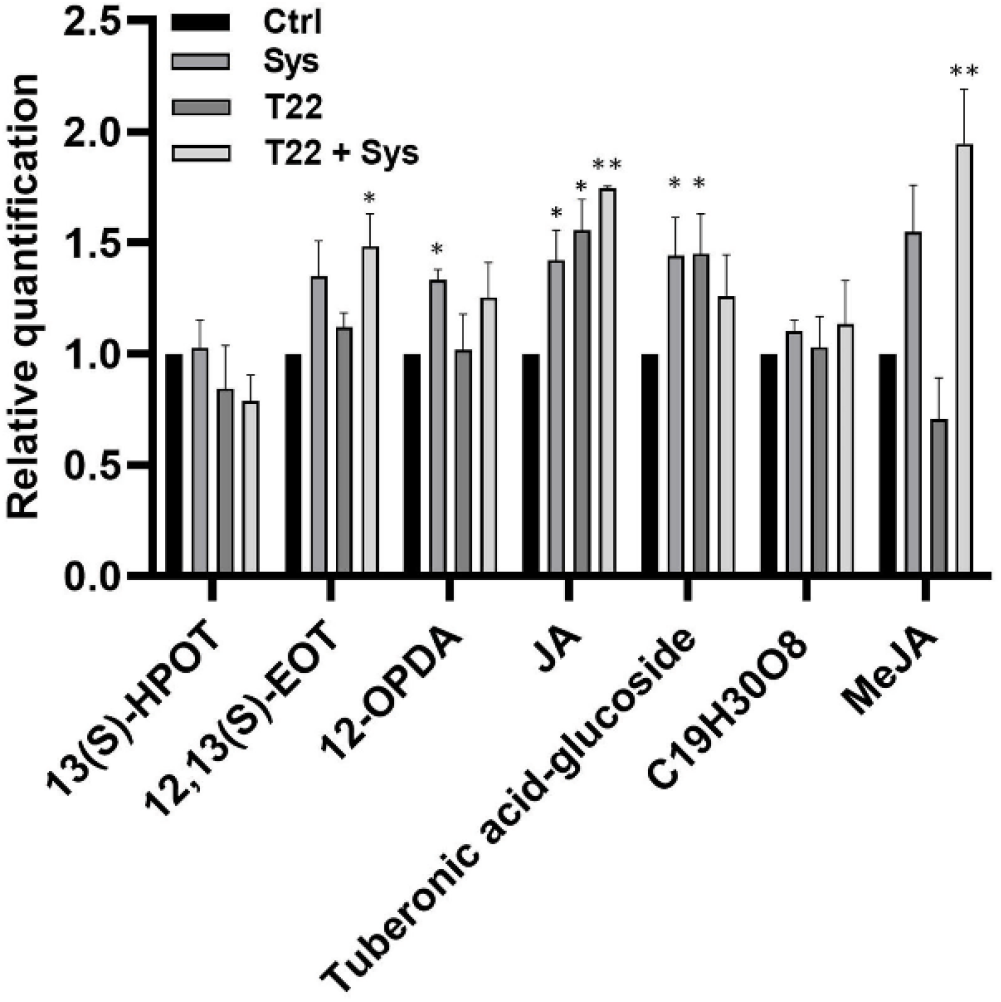
Targeted LC-HESI (−)-HRMS analysis of semi-polar metabolites in the JA pathway. Metabolites were relatively quantified as fold internal standard level, normalized on the control (Ctrl; for more details, see materials and methods). Data represents the average of three independent biological replicates, and a student’s t-test was used to identify metabolites whose abundance significantly changed with respect to the control condition, plants from control seeds treated with PBS 1X (* p ≤ 0.05; ** p ≤ 0.01).

Accumulation of tuberonic acid-glucoside, involved in JA metabolization, was higher in single (Sys and T22) treatments, and unaltered in T22+Sys plants (Figure 5). Finally, MeJA resulted over-accumulated in plants treated with both Sys and T22 (1,94 fold higher than control)

## 4 Discussion

The discovery of novel strategies for pathogen or pest control able to reduce the use of chemical pesticides represents an important challenge in modern agriculture. Beneficial microorganisms and natural-derived molecules with the potential to prime plant defenses are intriguing tools for the development of new generation biopesticides which could increase the sustainability of agricultural production. Some soil-dwelling fungi belonging to the genus *Trichoderma* have a profound impact on plant growth and protection limiting plant pathogen populations through mycoparasitism, competition and antibiosis (Harman et al., 2004; Verma et al., 2007; Yuan et al., 2019). In addition to pathogen control, root colonization by *Trichoderma* spp. limits pest-induced damage on plants either through stimulation of plant defense or by attracting natural enemies of insect pests (Tucci et al., 2011; Battaglia et al., 2013; Coppola et al., 2019b). Despite the efficacy of these fungal species in promoting plant growth and resistance against biotic threats, their potential can be limited in certain ecological contexts where their fitness is low (Di Lelio et al., 2021). To overcome this limitation, a combination of *Trichoderma* and other natural-derived elicitors of plants endogenous defense might represent an effective strategy to boost *Trichoderma* antagonistic ability both in fungal-friendly or harsh environments. Therefore, it seems feasible to combine the use of these root symbionts with natural-derived compounds, including plant-derived ones (Isman, 2006), semiochemicals (Witzgall et al., 2010), and interfering RNAs (Zhu et al., 2011; Koch et al., 2016) to attain satisfactory levels of plant protection. Natural molecules able to trigger plant immunity, such as Damage Associated Molecular Partners (DAMPs), are interesting players in pathogen and pest management. One of these is Sys, which acts as a DAMP in tomato plants following insect or mechanical damage by triggering plant immunity (Ryan, 2000; Ryan and Pearce, 2003). Coppola et al (Coppola et al., 2019c) and Molisso et al (Molisso et al., 2021) demonstrated that the exogenous supply of the peptide to intact healthy plants severely counteracted *S. littoralis* and *B. cinerea* fungal growth. These results are likely linked to the induction of plants defense-related genes through the accumulation of active compounds.

The absence of observed effects of Sys on T22 fitness was an important prerequisite to carry out the combined treatments. In fact, eliciting the same metabolic responses in the plant the two could have an antagonistic effect on each other. Indeed, *Trichoderma* and Sys ability to suppress different biotic threats resides in their propensity to prime plant defense responses through the modulation of different signaling pathways, including those of JA, ET and SA (Shoresh et al., 2005; Martínez‐Medina et al., 2017; Yuan et al., 2019). Priming is an intrinsic part of induced resistance during which the plant ‘projects’ a defense strategy against the potential invader while preparing its defensive approach for a faster and/or stronger reaction in future challenges (Mauch-Mani et al., 2017). The analysis of the expression of JA-associated genes in T22 and in Sys treated plants showed an up-regulation of *Aos*, an early defense-related gene, active in the octadecanoid signaling pathway that leads to JA biosynthesis, and of *Pin I* and *TD,* two late defense-related genes responsive to JA. It is well described that JA activates a cascade of defense responses that are temporally (early and late) and spatially (local and systemic) regulated in tomato (Ryan, 2000). The early and late wounding-response genes are regulated by distinct mechanisms. Local wounding leads to the processing of ProSys followed by Sys binding to SYR1, a membrane located receptor (Wang et al., 2018), and the subsequent activation of genes involved in JA biosynthesis. The increase of endogenous JA levels leads to the activation of late JA responsive genes that play a direct role in defense. Late defense genes include proteinase inhibitors (*PIs*), *TD*, and *LapA (Howe, 2004).* Tomato *LapA* is important in deterring herbivory-induced damage and controlling insect growth (Fowler et al., 2009), while *Pin I* prompts the formation of highly stable complexes with insect digestive proteases resulting in decreased digestion of dietary protein, depletion of essential amino acids, and, consequently, reduced rates of insect growth and development (Lorito et al., 1994). Furthermore, PIs in general and Pin I efficiently target microbial secreted proteases thus inhibiting a variety of plant pathogens including bacteria and fungi through suppression of cell growth (Hermosa et al., 2006; Turra and Lorito, 2011; Turrà et al., 2020). *TD* is a gene involved in the resistance against chewing insects. Remarkably, TD activity in the insect’s midgut is correlates with reduced levels of free threonine, which is a dietary requirement for phytophagous insects(Chen et al., 2007). We observed that the combination of the two treatments (Sys and T22) results in higher levels of transcripts related to the activation of the aforementioned genes, and of *LoxD,* a gene that also contributes to JA biosynthesis consequently, enhancing resistance against insect herbivores (Yan et al., 2013; Thakur and Udayashankar, 2019).

Metabolite data, obtained by LC-HRMS analyses, partly paralleled transcript profiles: single and double treatments were characterized by higher levels in JA, the end-product of the octadecanoic pathway involved in a series of physiological plant responses, in respect to control (Yang et al., 2019); the magnitude of alteration was stronger in T22+Sys plants compared to single treatments. A significant increase of MeJA, the JA volatile methyl ester, was also registered in plants simultaneously treated with both Sys and T22. Tuberonic acid-glucoside, a metabolite produced in the frame of JA catabolism, so called by virtue of its capacity to induce tuber formation in potato (*Solanum tuberosum*), also increased in Sys and T22 but not in the double-treated plants. Its synthesis, looks to be part of a general mechanism aimed to turning on and switching off JA levels and functions (Miersch et al., 2008): thus, in this context, the higher JA content in T22+Sys plants could reflect either the increased upstream metabolic flux, as highlighted at the gene expression level, or the reduced catabolism in the double compared to the single treatments. Additional increases in JA intermediates were found in Sys and T22+Sys samples, although this appears to be more related to stochastic events rather than to a solid tendency. The central role of JA in *Trichoderma* colonized plants in modulating defense responses has been demonstrated and confirmed through several mutant studies. The JA-deficient defenseless1 (def1) mutant plants, were found to be susceptible for disease development even after *Trichoderma* treatment, demonstrating that JA pathway is required for the resistance against pathogen challenged conditions (Martínez-Medina et al., 2013). It has also been shown that JA plays key roles during plant defense against necrotrophic fungal pathogens, such as *B. cinerea* and *Alternaria brassicicola*. The mutation of COI1, a receptor involved in JA signaling, makes plants more susceptible to *B. cinerea* (An et al., 2019). In this context, the stronger activation of the JA signaling pathway in Sys + T22 treated plants nicely correlates with the strong reduction of *B. cinerea* disease severity following the combined treatment compared to the single. In addition, we demonstrated that the combined treatment is highly effective against the soil-borne hemibiotrophic fungal pathogen *F. oxysporum*. Plants that had been simultaneously treated with both Sys and T22 showed a strong attenuation of mortality. Previous studies have shown that some *Trichoderma* spp. are able to inhibit or control the *Fusarium* wilt through the mechanism of mycoparasitism, production of lytic enzymes, and enhancement of the host endogenous defenses by the induction of plant hormones.(Mukherjee et al., 2012). It has been recently reported that JA is involved in *Trichoderma virens*-mediated resistance against *F. oxysporum* in tomato plants. The *T. virens* treatments on plants significantly reduced disease severity, increased JA levels and up-regulated the expression of JA-related genes (Jogaiah et al., 2018). A further recent study by Zehra et al (Zehra et al., 2017) demonstrated an integrated effect of *T. harzianum* and chemical inducers, SA and MeJA in tomato plants against *F. oxysporum* with increased expression of various antioxidant enzymes leading to elicitation of plant defense response.

Our findings showed for the first time that Sys strongly reduces the survival rate of larvae of *T. absoluta*, and corroborate previous findings showing that *Trichoderma* endophytic colonization negatively affects *T. absoluta* fitness through significant reduction of larval survival (Agbessenou et al., 2020). Furthermore, our study highlighted the usefulness of combined treatments to enhance plant resistance to *T. absoluta*. Previous works had shown a synergistic effect of different treatments on its fitness. For example, the combination of *B. bassiana* and *B. thuringiensis* strongly reduced *T. absoluta* larvae survival (Tsoulnara and Port, 2016). Yet, the usefulness of co-expression of different proteinase inhibitors to enhance plant resistance to *T. absoluta* has been highlighted (Hamza et al., 2018). This is the first study to evaluate the impact of peptide treatment and endophytic colonization.

Apparently, the combined application of Sys and T22 did not alter the oviposition behavior of *T. absoluta* with respect to the single powerful treatment with Sys. This result could have many explanations. For example, *T. absoluta* develops many overlapping generations/year, therefore mated females frequently oviposit on highly infested plants that are highly activated as Sys treatment. The combined treatment showed an additive effect on larval longevity/survival of *T. absoluta* in respect to the single treatments and this could correlate to the larger accumulation of JA-derived metabolites. It will be now interesting to test the effect of the combined treatments on the natural antagonists of *T. absoluta*, known to exploit plant VOCs to locate their victim (Lins et al., 2014; Ninkovic et al., 2021).

In conclusion, the combined application of *Trichoderma* and Sys on tomato plants enhances the level of JA and of its derivatives hence promoting a defense priming state able to protect treated plants by different biotic agents increasing the possibility to overcome the limits of the single agent usage. Since JA and its derivatives triggers VOCs emission (Tamogami et al., 2012), the JA increased level produced in treated plants may amplify the attractiveness of natural enemies of pests thus promoting indirect defenses of treated plants that could be protected also against other agents or biotic stresses.

## Author Contributions

AMA performed bioassays, gene expression studies; MC; supervised and participate to the experimental work; DT, SV designed and carried out experiments with *Fusarium oxysporum*, PC, EG designed and carried out *Tuta absoluta* bioassays; AF, GD, performed metabolomic analysis; CA, AR synthetized and purified Sys; VC contributed to gene expression analyses; SW supervised experimental work and revised the manuscript; RR conceived and supervised the work and wrote the manuscript. All the authors contributed to manuscript writing.

