## Additional file 1: Figure S1 for "Combination of the Systemin peptide with the beneficial fungus *Trichoderma afroharzianum* T22 improves plant defense responses against pests and diseases"

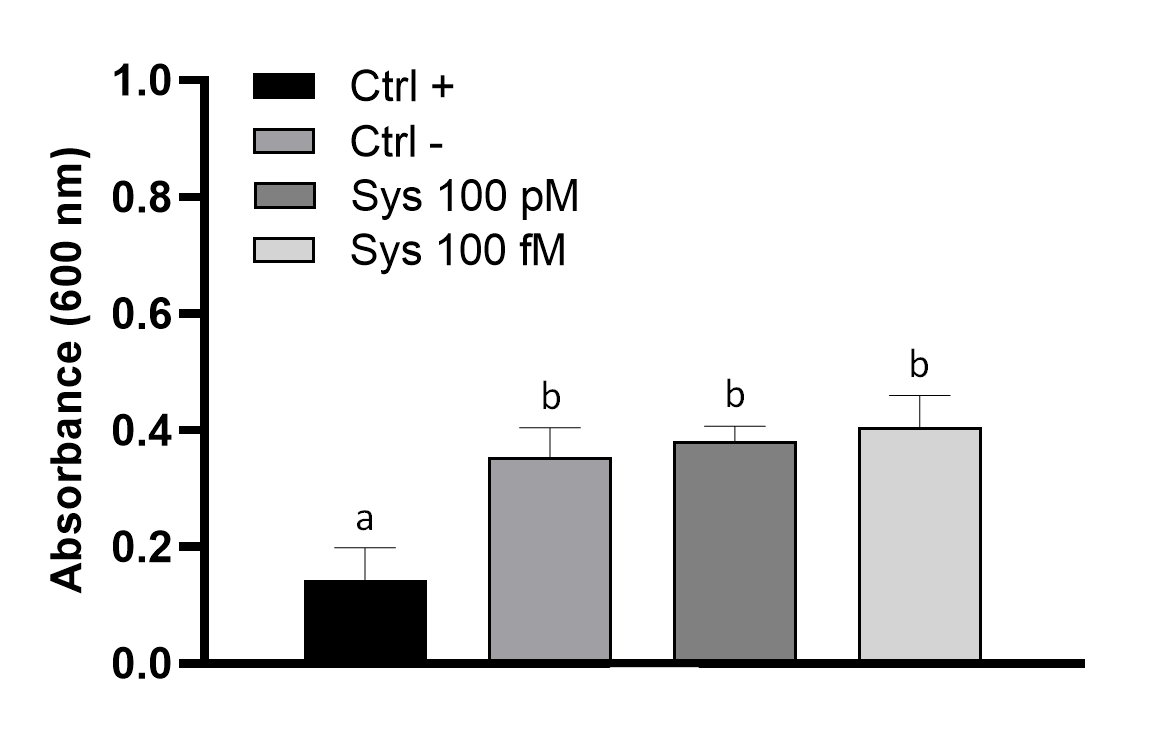

**Supplementary figure 1. Effect of Sys on *T. afroharzianum* T22.** Fungal growth in absence (Ctrl -) or presence of Sys (100 pM and 100 fM) was assessed 24h after fungal inoculation by assessing changes in optical density (OD_600_). Switch (200 µg/ml) was used as a positive control (Ctrl +). Different letters indicate statistically significant differences (One-Way ANOVA, P < 0.05). Error bars indicate standard error (n = 5).

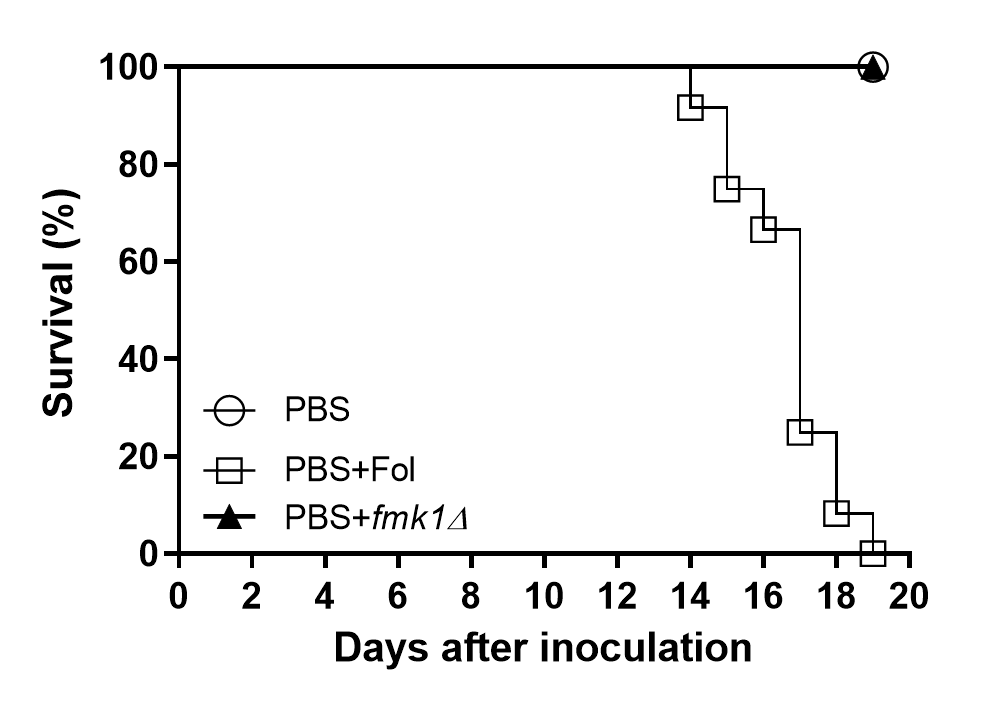

**Supplementary figure 2**. **Evaluation of *F. oxyporum* virulence on untreated tomato plants.** Kaplan–Meier plots showing survival of tomato plants germinated from PBS soaked seeds (Ctrl) and either left uninoculated or inoculated by dipping roots into a suspension of 5×10^6^ conidia/ml of the indicated *Fol* strains. Experiments were performed at least three times with similar results. Percentage survival of tomato plants was plotted for 20 days. Data shown are from one representative experiment.
