## Additional file 2: Table S1 for "Combination of the Systemin peptide with the beneficial fungus *Trichoderma afroharzianum* T22 improves plant defense responses against pests and diseases"

| **Table S1. List of Primers used for Real-time qPCR analysis.** | | |  |
| --- | --- | --- | --- |
| **Gene name** | **Primer** | **Sequence (5’-3’)** | **Accession number** |
| *AOS* | AOS Fw | GATCGGTTCGTCGGAGAAGAA | Solyc11g069800 |
|  | AOS Rv | GCGCACTGTTTATTCCCCACT |  |
| *EF-1α* | EF Fw | CTCCATTGGGTCGTTTTGCT | Solyc06g005060 |
|  | EF Rv | GGTCACCTTGGCACCAGTTG |  |
| *LoxD* | LoxD Fw | TTCATGGCCGTGGTTGACA | Solyc03g122340 |
|  | LoxD Rv | AACAATCTCTGCATCTCCGG |  |
| *Pin I* | Pin I Fw | GAAACTCTCATGGCACGAAAAG | Solyc09g084470 |
|  | Pin I Rv | CACCAATAAGTTCTGGCCACAT |  |
| *TD* | TrDeam Fw | TTAGACGCTTTCTCCCCTCGT | Solyc09g008670 |
|  | TrDeam Rv | GCTTGAGGAACTTGGAATCCC |  |
| *LapA* | Lap Fw | ATCTCAGGTTTCCTGGTGGAAGGA | Solyc12g010030 |
|  | Lap Rv | AGTTGCTATGGCAGAGGCAGAG |  |
